# Taxonomical and ontological analysis of verified natural and laboratory human coronavirus hosts

**DOI:** 10.1101/2023.02.05.527173

**Authors:** Yang Wang, Muhui Ye, Fengwei Zhang, Zachary Thomas Freeman, Hong Yu, Xianwei Ye, Yongqun He

**Author notes:** Co-Corresponding authors: Xianwei Ye, Yongqun He.

## Abstract

To fully understand COVID-19, it is critical to identify and analyze all the possible hosts of SARS-CoV-2 (the pathogen of COVID-19) and compare them with the hosts of other human coronaviruses. In this study, we collected, annotated, and performed taxonomical and ontological analysis of all the reported and verified hosts for all human coronaviruses including SARS-CoV, MERS-CoV, SARS-CoV-2, and four others that cause the common cold. A total of 37 natural hosts and 19 laboratory animal hosts of host human coronaviruses were identified based on experimental or clinical evidence. Our taxonomical ontology-based analysis found that all the verified susceptible natural and laboratory animals belong to therian mammals. Specifically, these 37 natural therian hosts include one wildlife marsupial mammal (i.e., Didelphis virginiana) and 36 Eutheria mammals (a.k.a. placental mammals). The 19 laboratory animal hosts are also classified as placental mammals. While several non-therian animals (including snake, housefly, zebrafish) were reported to be likely SARS-CoV-2 hosts, our analysis excluded them due to the lack of convincing evidence. Genetically modified mouse models with human Angiotensin-converting enzyme 2 (ACE2) or dipeptidyl peptidase-4 (DPP4) protein were more susceptible to virulent human coronaviruses with clear symptoms. Coronaviruses often became more virulent and adaptive in the mouse hosts after a series of viral passages in the mice. To support knowledge standardization and analysis, we have also represented the annotated host knowledge in the Coronavirus Infectious Disease Ontology (CIDO) and provided ways to automatically query the knowledge.

## Introduction

Zoonotic coronaviruses have caused dramatic impacts on humans. The existing COVID-19 pandemic has caused a disaster in public health worldwide. SARS-CoV-2, the cause of COVID-19, is a coronavirus that infects humans and leads to severe acute respiratory syndrome in humans. By the end of 2022, COVID-19 had caused over 660 million confirmed cases and over 6.6 million deaths worldwide. In addition to SARS-CoV-2, two other coronaviruses also caused major losses in this century. In 2002, Severe Acute Respiratory Syndrome (SARS) emerged in China, and eventually caused 8,098 confirmed human cases in 8 months and 774 deaths in 29 countries (1, 2). In 2012, the Middle East respiratory syndrome coronavirus (MERS-CoV) outbreaks, initially found in Saudi Arabia (3), resulted in 2,260 cases and 803 deaths across 27 countries (4, 5). In addition, four other human coronavirus strains, including HCoV-229E, HCoV-NL63, HCoV-OC43, and HCoV-HKU1, were also found worldwide and caused common cold in humans (6). It is critical to understand where coronaviruses originated and how they cause human outbreaks.

The human coronaviruses are likely able to spread and transmit from animals to humans. These coronaviruses appear to break down the species barrier through the transmission of natural host and likely intermediate host, and achieve the replication of the virus in the human body. Understanding how the virus spreads from host species to humans is crucial to preventing a pandemic. Previous studies have found many natural and laboratory animal hosts of human coronaviruses, such as bats (7), civets (8), camels (9), deer (10), monkeys (11) and mice (12). However, the exact scope of the human coronavirus hosts and their transmissional relations still remain unclear.

In the informatics domain, ontology is a structured vocabulary that represents entities and the relations among entities in a specific domain using a human- and computer-interpretable format (13). Many biological and biomedical ontologies have been developed and widely used. For example, the NCBITaxon ontology is a taxonomy ontology developed based on the classification of various types of cellular organisms and noncellular self-replicating organic structures including viruses in the NCBI taxonomy database (14). The Coronavirus Infectious Disease Ontology (CIDO) is a community-based ontology that systematically represents various coronavirus-related topics, including etiologies, hosts, transmissions, diagnosis, drugs, and preventions (15–17).

This study aims to survey and identify all verified hosts of human coronaviruses based on literature and database data collection, followed with systematic analysis of these human coronavirus hosts using ontology classification and bioinformatics methods. Given the current COVID-19 pandemic status, we have focused our study on the hosts of SARS-CoV-2. Our study found that all verified natural and laboratory animal hosts of human coronaviruses belong to Eutheria mammals (i.e., placental mammals). Genetically modified mouse models appeared to be more susceptible to virulent human coronavirus infections developing similar clinical symptoms, suggesting that host susceptibility depends on many factors including genetic modification for coronavirus binding to host cells.

## Methods

### Collection and annotation of verified hosts of human coronaviruses

Different resources were queried to identify verified animal hosts of human coronaviruses. Peer-reviewed journal articles from PubMed were mined and annotated to identify various hosts for different human coronaviruses, including natural hosts and laboratory animal models with experimental or clinical evidence. The PubMed searching keywords commonly used include: (SARS-CoV OR SARS-CoV-2 OR MERS-CoV OR “human coronovirus”) AND host. Meanwhile, the other resources including WHO reports and trusted newspapers are also searched. In addition to SARS-CoV, SARS-CoV-2, and MERS-CoV, there are also four human coronaviruses that cause common cold, which include HCoV-229E, HCoV-NL63, HCoV-HKU1 and HCoV-OC43. All these human coronaviruses are within our research scope. The reported evidence for being a host for a human coronavirus was then extracted, annotated, and recorded in a pre-designed Excel file. The recorded information was also summarized in formal tables and provided in the manuscript.

To be determined as a host of human coronaviruses, the required evidences are supposed to include at least one experimental confirmation using methods such as virus isolation, genomic sequencing, RT-PCR, and antibody neutralization assay. Clinical evidences such as related symptoms are recorded but not required for inclusion in our collection. This evidence is applied for wild type animals at its natural situation only, and transgenic animals are not counted. Although transgenic mouse model is much more effective model for human coronavirus study, wild type mice would also be infected with some human coronaviruses such as SARS-CoV-2 (B.1.351) (18) and MERS-CoV (19). Therefore, mouse is considered as a host. Zebrafish is a kind of vertebrate that share a high degree of sequence and functional homology with mammals. However, zebrafish is not included as a laboratory animal model because zebrafish has not been found effectively infectable with human coronaviruses probably due to dissimilarities between human and zebrafish ACE2 in the Spike-interaction region (20). Natural exposure or microinjection in different anatomic locations, including the coelom, pericardium, brain ventricle, or bloodstream, led a quick decrease of SARS-CoV-2 RNA in wild-type zebrafish larvae. After inoculated in the swim bladder (an aerial organ sharing similarities with the mammalian lung), the detected coronavirus decreased within 24 hours and then became stable through qRT-PCR, however, no clear evidence for production of new SARS-CoV-2 virions was observed (20), suggesting the failure of achieving detectable infection even in the swim bladder. A mosaic overexpression of hACE2 was not sufficient to achieve detectable infectivity of SARS-CoV-2 in zebrafish embryos or in zebrafish cells in vitro (20), further justifying the exclusion of zebrafish as a laboratory human coronavirus host. However, further study revealed that the humanized zebrafish, xeno-transplanted with human lung epithelial cells, could be a good model for SARS-CoV-2 infection (21).

If the required evidence as described above is not met, a suspected animal is not considered as host. For example, the Wikipedia contains a web page that lists animals that can get SARS-CoV-2 (22). After careful examination, three animals listed on the Wikipedia, including swan, zebrafish, and housefly, were not included in our verified list of SARS-CoV-2 hosts because the evidences provided for these animals could not be traced and verified. For instance, swan was cited in WHO report (23). However, a careful examination of the report and its cited resource could not find the required evidence of including swan as a host of SARS-CoV-2. Housefly, an insect, was also found to contain SARS-CoV-2 virus (24). However, it is very likely that the houseflies took up the blood fluids of the COVID-19 patients and virus RNA survived in the insect body without replicating, which means that the houseflies serve as a vector instead of host. This hypothesis was further confirmed by another independent study (25). Therefore, swan and housefly are not included in our host list.

An additional effort was also used to identify genetically modified mouse models used for study of human coronaviruses. Those mouse models susceptible to the infection of SARS-CoV, MERS-CoV and SARS-CoV-2 were particularly searched as these human coronaviruses are more important and have been deeply researched.

### Taxonomical classification of human coronaviruses and their hosts

To identify the specific taxonomical classification of human coronavirus and their host species, we extracted the hierarchical structure of various coronaviruses, and verified natural and laboratory animal hosts from the NCBI Taxonomy ontology (NCBITaxon) using the Ontofox tool (26), and the results were displayed using Protégé-OWL 5.5 editor (27). For Ontofox to run, the NCBI Taxonomy ontology IDs of specific species were used as the input. The Ontofox choice of “includeComputedIntermediates” was used to compute and retrieve the closest ancestors of different species in the hierarchical taxonomical tree. In addition to the different levels of ancestors, the Ontofox tool can also extract the relations among different levels of taxonomical terms and specific annotations such as scientific species names (i.e., labels) and common names (e.g., synonyms). The Ontofox output files were opened using the Protégé-OWL editor for visualization of the hierarchical structure and annotations. Screenshots of such visualization of human coronaviruses, nature hosts and laboratory models, which display the relationships among different taxonomical terms, were finally generated and saved as images.

### ACE2 Phylogenetic analysis method

A phylogenetic analysis was performed to establish the phylogenetic relations among different hosts of human coronaviruses using a method previously reported (28). Specifically, the ACE2 protein sequences from different host species of human coronaviruses were found from the NCBI Protein Database, and first aligned using the ClustalW program within the MEGA software (29). The phylogenetic tree of these proteins was then generated with the META tool using a Neighbor Joining method (30). To evaluate the robustness of identified phylogenetic tree structure, a Bootstrap analysis using 500 repetitions was implemented.

### CIDO ontological representation and analysis of human coronavirus-host relations

To support data standardization, integration, and analysis, we used the Coronavirus Infectious Disease Ontology (CIDO) (15, 16) to ontologically model, represent, and analyze the relations between human coronaviruses and hosts. The eXtensive Ontology Development (XOD) strategy (31) was applied for the CIDO ontology development. Specifically, new CIDO design patterns and axioms were first generated to semantically link different terms including the coronaviruses and their hosts. The Ontorat tool (32) was then used to transform an input data of human coronaviruses and their related hosts collected in Excel to OWL format for further display, editing, and analysis with Protégé-OWL 5.5 editor (33). To demonstrate the usage of the CIDO representation, the SPARQL RDF query language was used to query the Ontobee triple store that contains the CIDO knowledge represented as “subject-predicate-object” triples (34).

## Results

### Taxonomical classification of various human coronaviruses

Figure 1 shows the taxonomical classification of seven human coronaviruses, together with the Infectious bronchitis virus (IBV) (an avian coronavirus as control), and their relations under the hierarchy of taxonomy. Specifically, coronaviruses are positive stranded RNA viruses, belonging to the order Nidovirales, family Coronaviridae, and subfamily Coronavirinae. The subfamily Coronavirinae contains the four genera Alpha-, Beta-, Gamma-, and Deltacoronavirus. SARS-CoV and SARS-CoV-2 are members of the Sarbecovirus subgenus under the genus Betacoronavirus. MERS-CoV falls in Merbecovirus under the same genus Betacoronavirus (Figure 1) (35). The four coronaviruses that cause the common cold are also under the subfamily of Orthocoronavirinae. Specifically, HCoV-229E is the member of the Duvinacovirus subgenus, HCoV-NL63 is the member of the Setracovirus subgenus, and they are both under the Alphacoronavirus genus. HCoV-HKU1 and HCoV-OC43 are members of the Embecovirus subgenus under the genus Betacoronavirus. The IBV avian coronavirus, which does not infect humans, is a member of the Gammacoronovirus (Figure 1).

**Figure 1.**
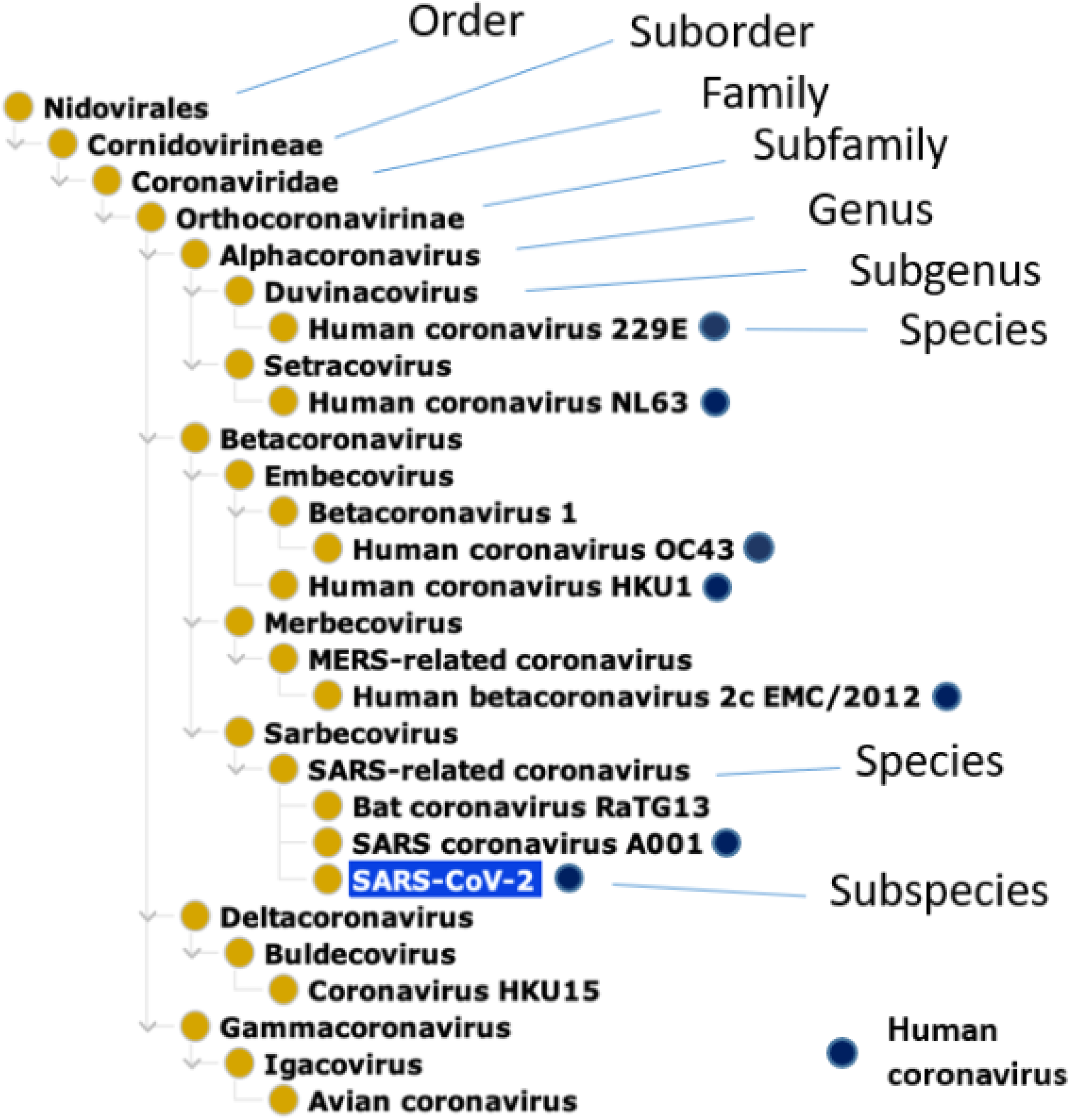
Taxonomical hierarchy of human coronaviruses based on the NCBITaxon Ontology. The dark blue circles label those coronaviruses capable of infecting human. Under the subgenus Sarbecovirus and genus Betacoronavirus, SARS–related coronavirus is a species that includes specific SARS-CoV-1 and SARS-CoV-2 strains. Given that there are many specific strains of MERS-CoV and SARS-CoV speces, only representative strains are shown here. Bat coronavirus RaTG13 (a bat coronavirus strain highly homologous to SARS-CoV-2), Coronavirus HKU15 (a coronavirus species that infects pigs), and Avian coronavirus (a coronavirus species that infects birds) are also included here as examples of non-human coronaviruses.

The taxonomical classification of the coronaviruses appears to be associated with the host species that these coronaviruses turn to infect. In general, Alpha- and Betacoronaviruses mainly infect mammalian species including humans, and Gamma- and Deltacoronaviruses primarily infect birds (6). Note that although bats can fly, it is a mammalian species. Bat-borne betacoronaviruses have been found to be closely related and responsible for many human respiratory infections (36).

### Identification and classification of 37 verified natural animal hosts of human coronaviruses

In this article, natural hosts of human coronaviruses are defined as those animals identified to be infected with human coronaviruses with experimental evidence. Table 1 collects 37 natural hosts of different types of human coronaviruses based on convincing evidence as reported in the literature. The natural hosts infected with SARS-CoV-2 with solid evidence include tiger, cat, dog, pangolin, lion, gorilla, and white-tailed deer (Table 1).

**Table 1.** Verified natural hosts of human coronaviruses (in alphabetic order)

| Host | Taxon ID | Human Coronaviruses | Evidence | References (PMID or reference citation) |
| --- | --- | --- | --- | --- |
| Human | 9606 | SC, SC2, MC, OC43, HKU1, 229E, NL63 | Virus isolation, RT-PCR, serum antibody detection, symptoms. | 15347429, 26695637, 29551135, 32798769 |
| Alpaca | 9655 | MC | Seroepidemiological study | 30832356 |
| Asian small-clawed otter | 452597 | SC2 | PCR, Mild respiratory symptoms including sneezing, runny noses, mild lethargy, and coughing | (77) |
| Bactrian camel | 9837 | MC | Antibody detection, virus isolation | 31969478 |
| Bat (e.g., Chinese rufous horseshoe bat) | 9397 (e.g., 89399) | SC, MC, 229E, NL63 | Genome sequencing | 30844511, 16195424, 30531947, 26695637, 29551135 |
| Binturong | 94180 | SC2 | PCR and sequencing | 36121159 |
| Black-Tailed Marmoset | 1090896 | SC2 | Viral RNA detection, immunohistochemistry detection, | 35577455 |
| Canada Lynx | 61383 | SC2 | Respiratory symptoms, RT-PCR | 36121159 |
| Cattle | 9913 | OC43 | Genetic sequencing | 32629960,15650185 |
| Chinese ferret-badger | 204267 | SC | Neutralizing antibody detection, RNA isolation, RT-PCR | 12958366 |
| Civet | 9673 | SC | PCR, virus isolation, neutralizing antibody detection | 30844511 |
| Coatimundi | 743424 | SC2 | PCR and sequencing | (78) |
| Cougar | 9696 | SC2 | PT-PCR, anorexia, diarrhoea, and nasal discharges | 36121159, 35062324 |
| Dog | 9615 | SC2 | Genomic sequencing, virus isolation | 32408337 |
| Domestic cat | 9685 | SC, SC2 | ELISA, neutralization assay RT-PCR | 15921605, 32402157 |
| Domestic mouse | 10090 | HKU1 | PCR | 19239338 |
| Dromedary camel | 9838 | MC, 229E | Antibody detection, virus isolation | 31969478, 24896817, 27528677 |
| Fishing cat | 61388 | SC2 | Lethargy and gastrointestinal signs, PCR and sequencing | 36121159 |
| Giant anteater | 71006 | SC2 | Viral RNA detected in rectal swab, spleen and kidney | 36692797 |
| Gorilla | 9593 | SC2 | Nuclear sequence detection | (79) |
| Hippopotamus | 9833 | SC2 | PCR and coughup some snot | (80) |
| Indian Leopard | 421001 | SC2 | Whole-genome sequence | 36121159 |
| Large hairy armadillo | 29080 | SC2 | Respiratory symptoms, RT-PCR | (81) |
| Lesser rice-field rat | 69075 | SC | RT-PCR | 15921605 |
| Lion | 9689 | SC2 | Virus detection | 33051368 |
| Malayan pangolin | 9974 | SC, SC2 | Metagenomics prediction | 31652964, 32197085 |
| Masked palm civet | 9675 | SC | PCR, virus isolation, neutralizing antibody detection | 12958366, 23671097 |
| Mink | 9655 | SC2 | Genomic sequencing, PCR | 33172935, 32663073 |
| Pangolin | 9974 | SC2 | Genomic sequencing | 32724171 |
| Raccoon dog | 34880 | SC | RT-PCR, virus isolation, neutralizing antibody detection | 12958366 |
| Red fox | 9627 | SC | RT-PCR | 16485471 |
| Snow Leopard | 29064 | SC2 | Mild respiratory signs, occasional dry cough, wheezing, PCR confirmation | 36121159 |
| Spotted hyenas | 9678 | SC2 | Mild lethargy, nasal discharge, and occasional cough, PCR | 36121159 |
| Tiger | 9694 | SC2 | Virus detection | 33051368 |
| Virginia opossum | 9267 | SC2 | Nasopharyngeal (NP) swabs and sera samples, PCR | (82) |
| White-footed mouse | 10041 | SC2 | Nasopharyngeal (NP) swab RT-PCR, and serum antibody detection. | (82) |
| White-tail deer | 9874 | SC2 | Neutralization test, rPT-PCR | 34942632, 3578920 |

Based on the hierarchical classification, all the 37 verified natural hosts of human coronaviruses are categorized under the Theria clade. These 37 therian mammals can be classified to a wildlife marsupial mammal (i.e., *Didelphis virginiana*) and 36 Eutheria mammals (a.k.a. placental mammals) (Figure 2). *Didelphis virginiana* is also called Virginia opossum, which belongs to Didelphimorphia order under Metatheria clade. Metatherian mammals, also known as marsupials, are an ancient group, and most marsupials are found in Australasia (around 200 species) and Central and South America (around 70 species). A study conducted by Goldberg et al. found 8% of wild life Virginia opossum having detectable serum antibody against SARS-CoV-2 (37).

**Figure 2.**
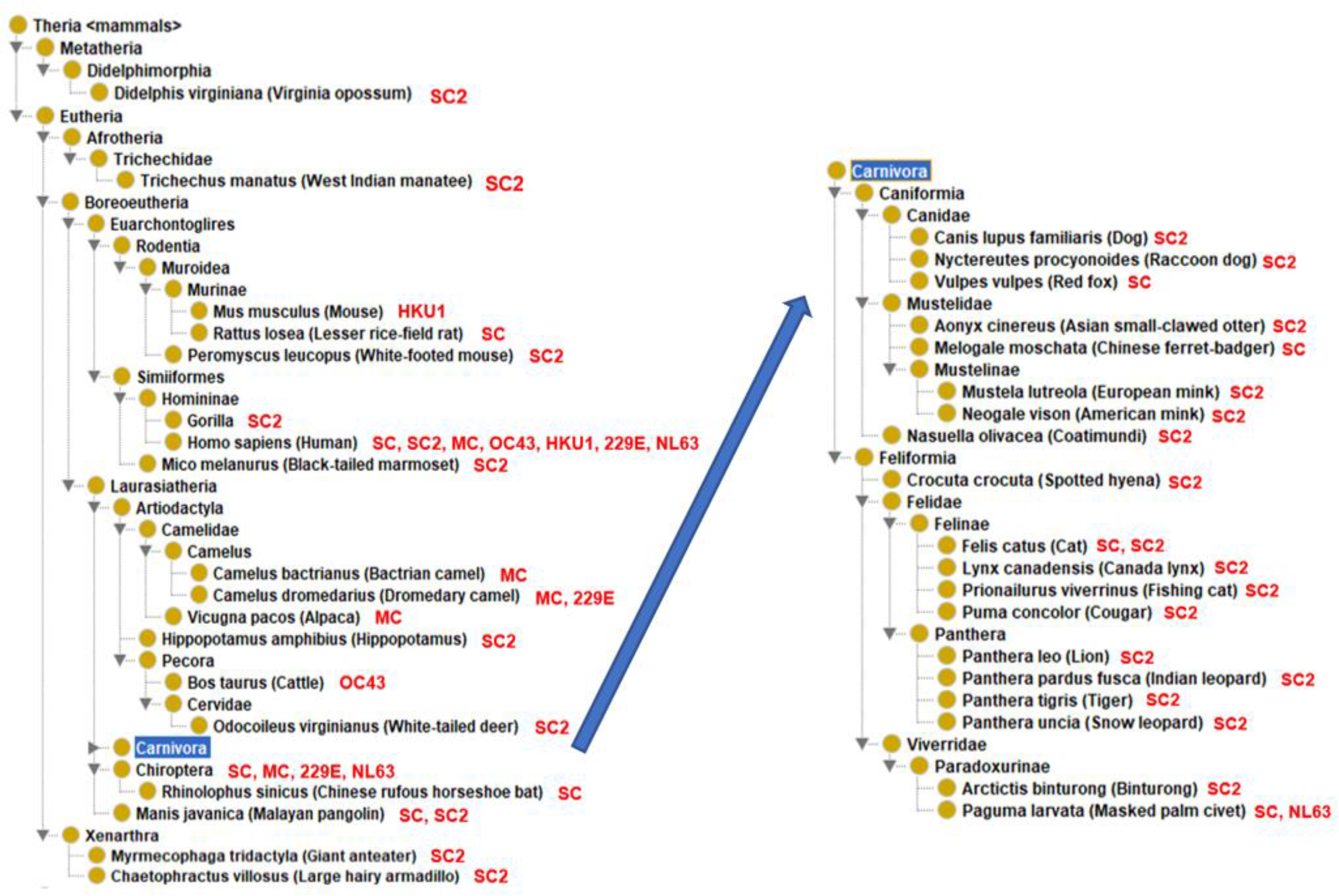
Taxonomical hierarchy of 37 verified natural hosts of human coronaviruses. All these hosts belong to Theria mammals. The hierarchy of the Carnivora animal hosts is singled out for the purpose of optimal visualization. The added red-color name abbreviations indicate the coronaviruses capable of infecting the corresponding animal host. Abbreviations: SARS-CoV-2: SC2; SARS-CoV: SC; MERS-CoV: MC; HCoV-OC43: OC43; HCoV-HKU1: HKU1; HCoV-229E:229E; HCoV-NL63: NL63.

Eutheria is synonymous with “placental mammals” and includes placental mammals and their closest extinct relative (38). In the taxonomical classification, most of our verified natural hosts are categorized within the superorders Euarchontoglires and Laurasiatheria under the clade Boreoeutheria, and only two (i.e., large hairy armadillo and giant anteater) under the superorder Xenarthra. Euarchontologilres include Homininae and Murinae (rodents) subfamilies (Figure 2). The Homininae subfamily contains *Homo sapiens* (human) and gorilla. In addition to humans, the world’s first positive case of gorillas infected with COVID-19 was found in a California zoo (39). In rodents, HCoV-HKU1 is currently considered to be a rodent-related virus, originally obtained from infected mice (Figure 2) (40). In 31 animals sampled in January 5, 2004 before culling of wild animals at a Guangzhou live animal market, including 4 cats, 3 red fox and one Lesser rice field rats were tested SARS-CoV positive based on an RT-PCR test (41).

Many natural hosts are classified under Laurasiatheria (Figure 2). Classified under the Rhinolophus genus of the order Chiroptera, bats have been found to be the host of SARS-CoV, MERS-CoV, HCoV-229E, and HCoV-NL63 based on the isolation of these viruses from bats and genomic sequencing confirmation (Figure 2, Table 1). SARS-CoV is known to exist in greater horseshoe bat and Chinese rufous horseshoe bat (Table 1). SARS-CoV-2 genome shows high homology to SARS-related coronaviruses identified in horseshoe bats (42). The sequence homology between SARS-CoV-2 and SARS-CoV is 79.6% (42). RaTG13, a bat coronavirus that shares 96% genetic similarity with SARS-CoV-2, was isolated from horseshoe bat (42). However, the exact SARS-CoV-2 virus or its genome sequence has not been isolated from bats through natural infections.

Malayan pangolin (*Manis javanica*) is a species under Laurasiatheria. Viral metagenomics showed that several Malayan Pangolins host a variety of coronaviruses, among which SARS-CoV was the most widespread one (43). Zhang et al. found a SARS-CoV-2-like CoV (named Pangolin-CoV) in dead Malayan pangolins, Pangolin-CoV is 91.02% and 90.55% identical to SARS-CoV-2 and BatCoV RaTG13, and concluded that except for RaTG13, Pangolin-CoV is the most closely related coronavirus to SARS-CoV-2 (44). On March 26, 2020, Yi Guan (45) detected several coronaviruses in a small number of pangolins smuggled into China that are closely related to SARS-CoV-2. However, this similarity is still insufficient to indicate that Malayan pangolins are intermediate hosts directly involved in the current SARS-CoV-2 outbreak.

Cattle, camel, and alpacas are classified under the Artiodactyla order (Figure 2). Camels, including Bactrian camels, dromedaries and hybrid camels, have been found to be an important intermediate host for MERS-CoV (46). MERS infection has also been found in alpacas that live with camels (47). HCoV-229E was also isolated from alpacas raised in captivity with dromedary camels (6). Using gene sequencing technology, Vijgen et al. found HCoV-OC43 and bovine coronavirus (BCoV) have remarkable antigenic and genetic similarities, and BCoV and HCoV-OC43 had a relatively recent zoonotic transmission event with their common ancestor likely dated to around 1890 (48).

The largest number of animal species found to host human coronaviruses are under the carnivora order (Figure 2). Under the Canidae family, dog, raccoon dog, red fox, Chinese ferret-badger, and mink were found to host SARS-CoV or SARS-CoV-2 (Figure 2, Table 1). The raccoon dog was first identified in 2003 as an intermediate host of SARS in addition to civets (41). Pet dogs were first reported to be infected with SARS-CoV-2 in Hong Kong and then reported subsequently in other places around the world (49). SARS-CoV-2 was isolated from minks (under Mustelidae) on farms in the Netherlands, leading to mass culling (50). Classified under the Feliformia suborder, domestic cat, lion, tiger, and civets were also found to be infected with human coronaviruses (Figure 2, Table 1).

### Identification and classification of 19 verified laboratory animal hosts of human coronaviruses

Laboratory animal models have been widely explored to study human coronaviruses with host-virus interaction mechanism and translational drug/vaccine studies. Our literature mining identified 19 laboratory animal models that have been used in various laboratory studies on human coronavirus hosts (Table 2). Human coronaviruses have been detected in these laboratory animals using experimental methods from different anatomical locations such as saliva, blood, and lungs, and many of the infected animals can develop symptoms and replicate in vivo (Table 2).

**Table 2.** Laboratory models of human coronavirus hosts (in alphabetic order)

| Host | Taxon ID | Human Coronaviruses | Evidence | References (PMID) |
| --- | --- | --- | --- | --- |
| African green monkey | 9534 | SC | Moderate to high titers of SARS-CoV, interstitial pneumonitis | 15527829 |
| Alpaca | 30538 | MC | Nasal swab specimens, serum samples and mild respiratory disease | 27070385 |
| Baboon | 9557 | SC2 | Detect viral RNA in nasopharyngeal swabs, and inflammation | 33340034 |
| Bank vole | 447135 | SC2 | qRT-PCR | 33754987 |
| Civet | 9673 | SC | RT-PCR | 17037579 |
| Common Tree Shrew | 9395 | SC2 | Elevated body temperature, virus detection | 32994418 |
| Common marmoset | 9483 | SC | Immunohistochemistry, RT-PCR | 16049331 |
|  |  | MC | Respiratory disease, qRT-PCR | 25144235 |
| Cynomolgus macaque | 9541/36519 | SC | Nasal, oral and rectal swabs, RT-PCR | 12748632, 21533129 |
|  |  | MC | Antibodies detection in sera, RT-qPCR | 32303590 |
|  |  | SC2 | Antibodies detection in sera, RT-qPCR | 32303590 |
| Ferret | 9669 | SC | Infected intranasally with 10(3) TCID<sub>50</sub> SARS-CoV | 18234270 |
|  |  | SC2 | Fever and inappetence, viral detection | 32269068 |
| Fruit bat | 9407 | SC2 | RT-qPCR, antibody detection | 32838346 |
| Mouse | 10090 | SC | Prior infection, neutralizing antibody detection | 15016880 |
|  |  | MC | Serum neutralizing antibodies and MERS-CoV S1 protein-specific IgG antibodies detection, even death | 26446606 |
|  |  | SC2 | expressing hACE2 receptors, PT-PCR | 32380511, 36222118 |
| North American raccoon | 9654 | SC2 | Intranasally inoculated, rPT-PCR, seroconverted | 35097038 |
| North American deer mouse | 10042 | SC2 | Virus detection in nasal washes, oropharyngeal and rectal swabs | 34127676 |
| Raccoon dog | 34880 | SC2 | Viral replication and tissue lesion in nasal conchae | 33089771 |
| Rhesus monkey | 9544 | SC | RT-PCR, virus isolation | 15892035 |
|  |  | MC | qRT-PCR | 24218506 |
|  |  | SC2 | Changes in respiratory pattern, piloerection, reduced appetite, hunched posture, dehydration, virus detection | 32396922 |
| Rabbit | 9986 | SC2 | qRT-PCR, viral RNA positive in the nose and throat for at least four days | 33356979 |
| Sheep | 9940 | SC2 | RT-qPCR of respiratory tract tissues and lymphoid tissues | 34816258 |
| Syrian hamster | 10036 | SC | Virus detection | 17037579, 17499378 |
|  |  | SC2 | Tachypnea, weight loss, inflammation, bronchopneumonia, virus detection | 32215622, 36222118 |
| Striped skunk | 30548 | SC2 | rPT-PCR, seroconverted | 35091038 |

**Abbreviations:** SARS-CoV-2: SC2; SARS-CoV: SC; MERS-CoV: MC.

Similar to the natural hosts of human coronaviruses as described above, all the 19 laboratory animal models belong to Boreoeutheria, a clade (magnorder) of Eutheria (i.e., placental mammals). More specifically, these laboratory animals are categorized under two clades of Boreoeutheria: Euarrchontoglires and Laurasiatheria (Figure 3). Different from the natural hosts, the list of existing laboratory animals does not include any species under the superorder Xenarthra.

**Figure 3.**
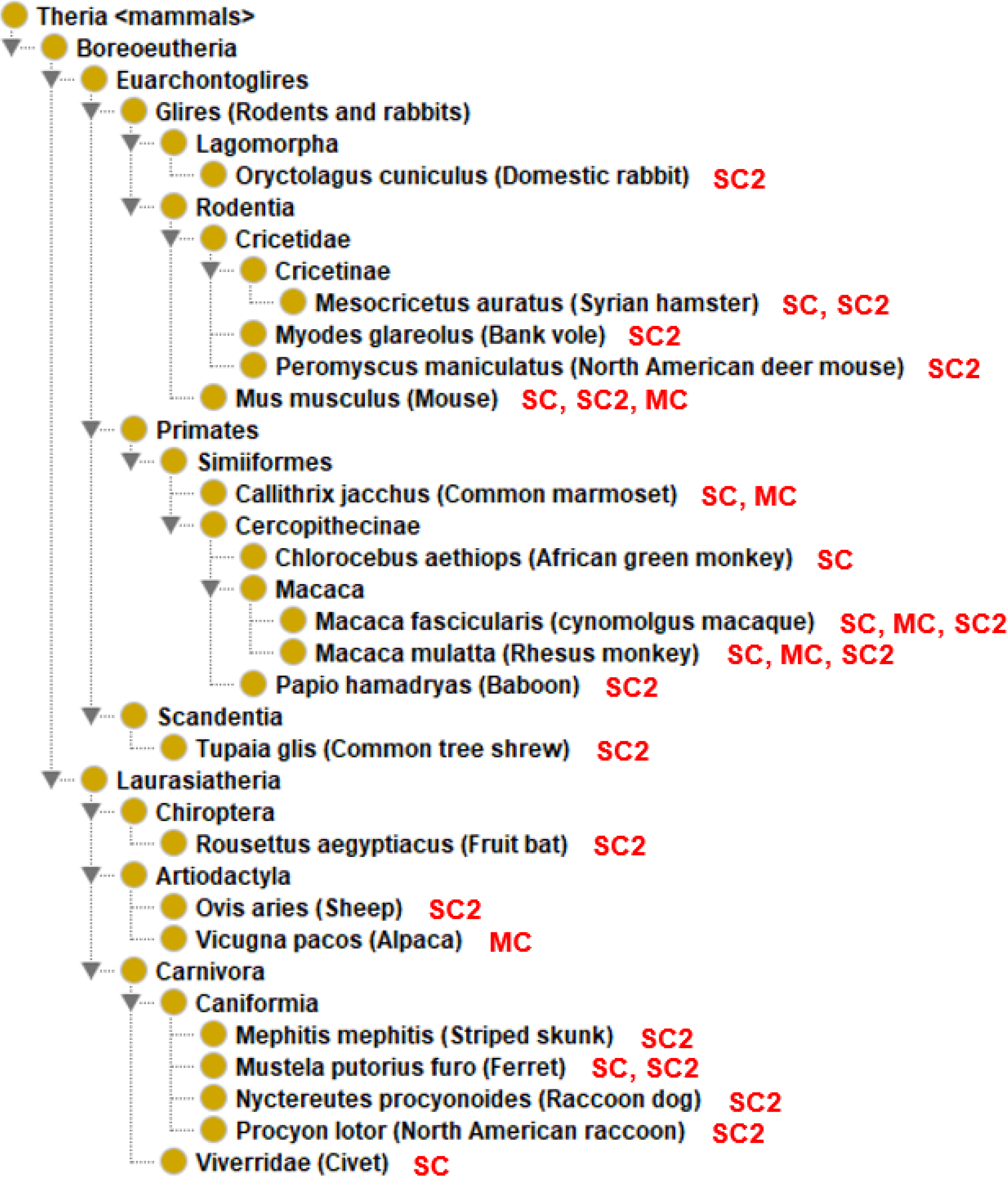
Taxonomical hierarchy of 19 laboratory animal hosts of human coronaviruses. All these hosts belong to Boreoeutheria, a special group under mammals. Abbreviations: SARS-CoV-2: SC2; SARS-CoV: SC; MERS-CoV: MC.

Ten laboratory animals are categorized under four orders Primates, Rodentia, Lagomorpha, and Scandentia within the Euarrchontoglires clade. Primates is the order of animals phylogenetically close to humans and ideal laboratory animal models for human coronaviruses (51–53). These laboratory animals under Primates are all Simiiformes (infraorder), including marmoset, African green monkey, and macaque. These non-human primates are frequently used for COVID-19 research; however, the usage of these animals is also expensive (54).

The orders Rodentia and Lagomorpha belong to the clade Glires. Under Glires are four laboratory animal species, including Syrian hamster, bank vole, deer mouse, and domestic mouse, and domestic rabbits (Figure 3). The first four belong to Rodentia and the last one belongs to Lagomorpha. The Lagomorpha have a second pair of incisor teeth behind the first pair in the upper jaw, while Rodentia have only one pair. The mouse is the most commonly used animal model for coronavirus research (55, 56). Mouse models are popular because of their affordability, availability, and clear genetic backgrounds, and they have been widely used for studying pathogenesis of human coronaviruses (57). Mouse has been found to be a natural host for human coronavirus strain HKU1 (Figure 2) (58). However, mice infected naturally with SARS-CoV-2 show fewer symptoms and low virus replication (59). To make mouse a better laboratory model for coronavirus studies, genetically modified mice are often used as detailed in the next section.

Under the Laurasiatheria clade are seven animals separated into two orders: Artiodactyla and Carnivora. Sheep and alpaca are under Artiodactyla. The five species under Carnivora include striped skunk, domestic ferret, raccoon dogs, and North American Raccoon (Table 2, Figure 3). Compared to the large number of Laurasiatheria animals in the group of natural human coronavirus hosts, the number of Laurasiatheria animals in the laboratory host group is relatively smaller, likely due to their expensive cost and less related reagents available for deep investigations.

### Various transgenic mouse models are developed for human coronavirus studies

To increase the infection rate, different transgenic mice were developed and utilized (55). A comparative study showed that genetically modified mouse model is better than the wild-type mouse model with adenovirus delivered hACE2 (59), suggesting that genetically modified mouse models are preferred model for coronavirus studies. The commonly used genetically modified mouse models for SARS-CoV, MERS-CoV and SARS-CoV-2 were collected and provided in **Table 3**.

**Table 3.**
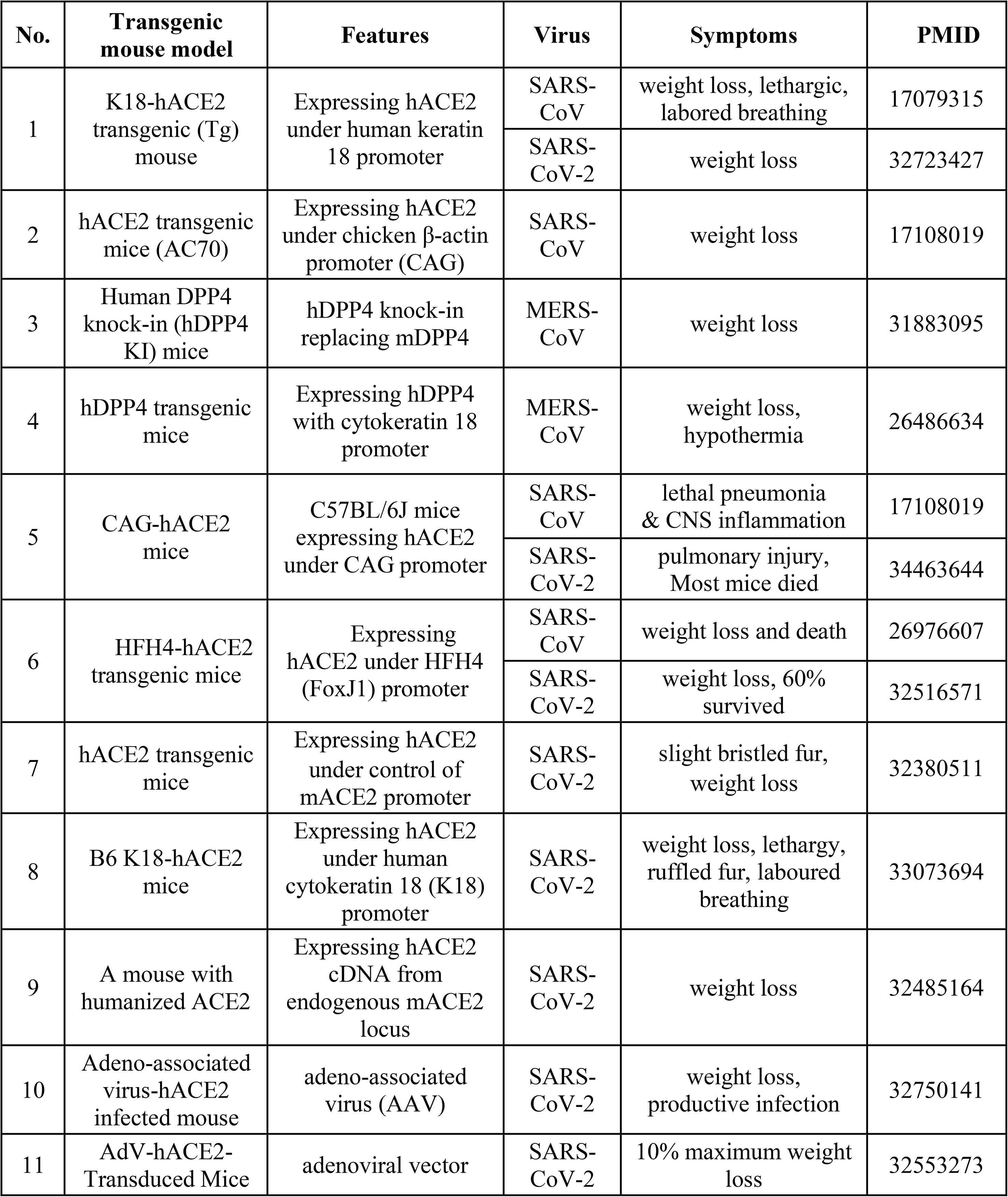
Genetically modified mouse models for SARS-CoV, MERS-CoV, SARS-CoV-2.

Typically, transgenic mouse models were typically generated to express hACE2 (human angiotensin-converting enzyme 2) or hDPP4 (dipeptidyl peptidase-4). ACE2 is the host receptor that binds the S protein in SARS-CoV and SARS-CoV-2, and the DPP4 is the receptor binding the S protein in MERS-CoV (60). An example of the genetically modified mouse model is the HFH4-hACE2 C3B6 mouse that expresses human ACE2 under the control of a lung ciliated epithelial cell-specific HFH4/ FOXJ1 promoter (61, 62). HFH4-hACE2 mice expressed high levels of hACE2 in the lung, but at varying expression levels in other tissues, including the brain, liver, kidney, and gastrointestinal tract (63).

Methods for generating hACE2 or hDPP4 mice differ (Table 3). Different promoters are used to drive the expression of the hACE2 or hDPP4 gene in mice. For example, the most commonly used mouse models of SARS-CoV and SARS-CoV-2 are transgenic mice with human cytokeratin 18 as the promoter and human ACE2 added (54). In addition, the chicken-β actin promoter (for SARS-CoV infection) and HFH4/ Foxji promoter (for SARS-CoV-2 infection) were also used to generate transgenic mice. The CRISPR/Cas9 knock-in technology has also been used to generate a mouse model expressing human ACE2 (hACE2) (64).

Genetically modified mice show significantly higher coronavirus infection rates and more severe symptoms than wild type mice. For example, while wild-type C57BL/6 mice showed no or low viral loads after intranasal infection with SARS-CoV-2, young and aged genetically modified hACE2 mice sustained high viral loads in lung, trachea, and brain (64). Although SARS-CoV-2 infected-aged hACE2 mice survived, interstitial pneumonia and elevated cytokines were observed. It was also found that intragastric inoculation of SARS-CoV-2 caused viral infection and pulmonary pathological changes in hACE2 mice (64).

To further improve the laboratory mouse model for coronavirus studies, continuous virus passaging of coronaviruses in wild type and even genetically modified mice was frequently used to generate mouse-adapted coronaviruses (17, 65). For example, a clinical isolate of SARS-CoV-2 strain was serially passaged for 6 generations in the respiratory tract of aged BALB/c mice, resulting in the generation of a more infectious genetically modified strain called MASCp6 (66). Adaptive mutations, including a N501Y mutation in the spike protein receptor binding domain, were later identified by deep sequencing of the MASCp6 genome (66). Similarly, Roberts et al. (65) generated the mouse-adapted SARS-CoV-2 MA15 strain after 15 passages of SARS-CoV-2 in BALB/c mice, and the resulting MA15 became lethal for mice following intranasal inoculation. Li and McCray utilized human DPP4 knock-in (hDPP4 KI) mice to infect MERS-CoV, but the transgenic mice still did not display respiratory disease after MERS-CoV infection. After serially passages of 30 generations in vivo, the wild type MERS-CoV strain eventually became MERS_MA_6.1.2, which produced significantly higher titers than the parental virus strain in the lungs of hDPP4 KI mice and caused diffuse lung injury and a fatal respiratory infection (67).

### Ontological modeling, query, and analysis of human coronavirus hosts

To support standardized and digitized modeling and representation of human coronavirus hosts, we represented our collected human coronavirus hosts in the Coronavirus Infectious Disease Ontology (15, 16). To represent the relations between coronavirus and its host, in CIDO we generated a new ontology relation called ‘*capable of infecting host*’, which represents a relation between a pathogen and a host in which the pathogen has verified evidence of infecting the host. For example, the following axiom relation defined in CIDO represents that SARS-CoV-2 is capable of infecting white-tail deer:

*SARS-CoV-2: ‘capable of infecting host’ some ‘white-tail deer’*

Using such strategy, we have represented all the human coronaviruses and their infected hosts as identified in Tables 1–2 and Supplemental File 1.

For genetically modified mouse models (Table 3), since they are not represented in NCBITaxon ontology or NCBI Taxonomy database, we have generated new terms of these specific genetically modified mouse terms in CIDO and developed new logic axioms to represent their properties. For example, the CAG-hACE2 mouse model expresses human ACE2 gene, which is ontologically represented as follows:

*CAG-hACE2 mouse: expresses some ‘angiotensin-converting enzyme 2 (human)’*

To demonstrate the usage of our ontological modeling, we provide a SPARQL program to demonstrate its usage for advanced query. Figure 4 provides a demonstration of a SPARQL script used to find the number of verified organisms that are capable of being infected by SARS-CoV-2 virus through SPARQL query of knowledge stored in CIDO. Specifically, 38 hits were identified from the SPARQL script execution. Supplemental File 2 provides another SPARQL query that lays out the specific details of these 38 SARS-CoV-2 hosts.

**Figure 4.**
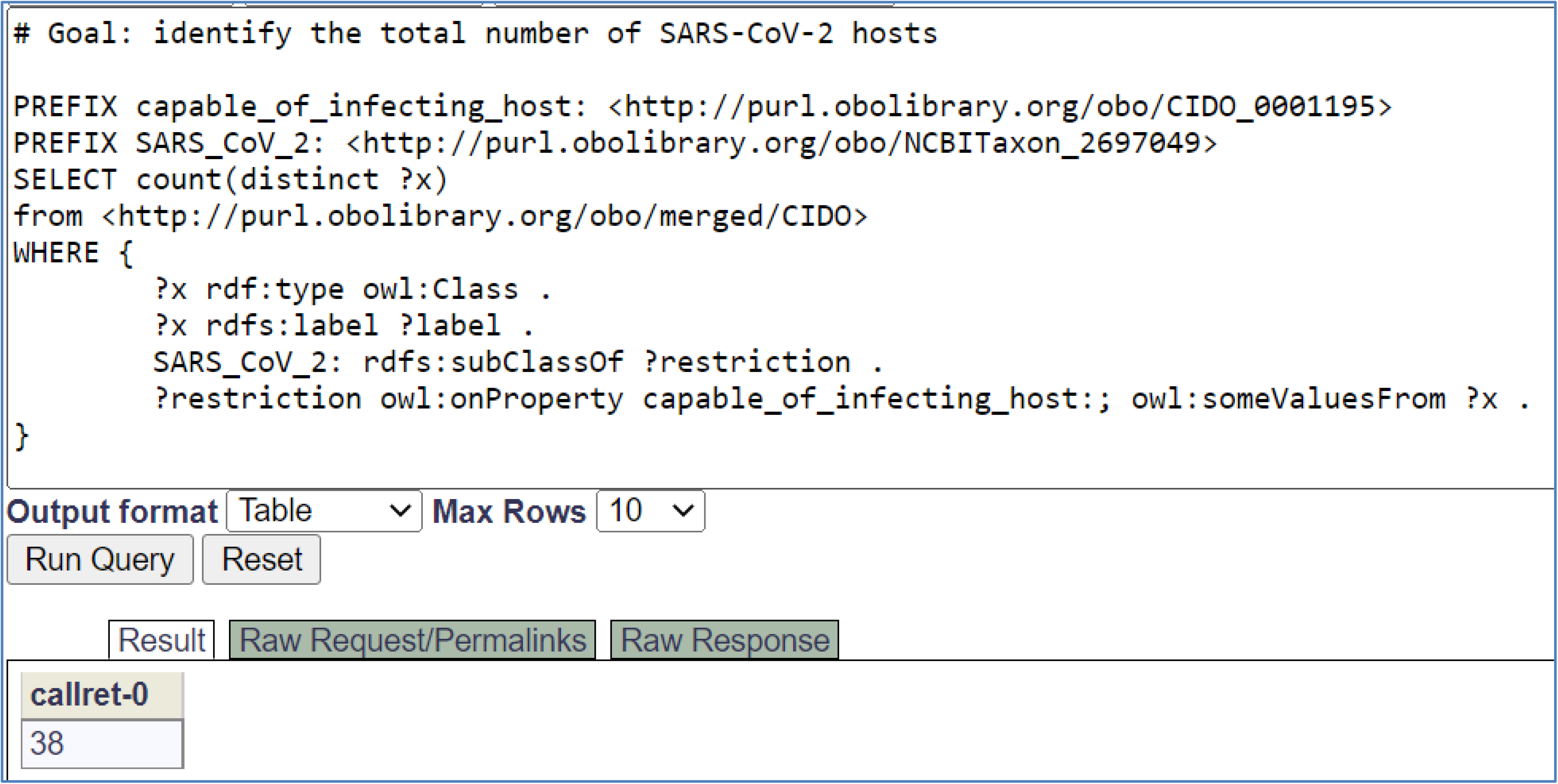
SPARQL query for SARS-CoV-2 hosts stored in CIDO. This SPARQL identified 37 organisms that are capable of being infected by SARS-CoV-2 virus. The SPARQL was performed using Ontobee SPARQL endpoint (http://www.ontobee.org/sparql). Another SPARQL query script that lists the names of these 38 hosts is provided in Supplemental File 2. The detailed information about these 38 hosts is also provided in Supplemental File 1, which offers the Excel sheets of the detailed hosts for different human coronaviruses.

## Discussion

The contributions of this article are multiple. First, we systematically surveyed, identified, and ontologically classified 37 natural and 19 laboratory animals of human coronaviruses (including SARS-CoV, SARS-CoV-2, MERS-CoV, and four non-virulent human coronaviruses) with experimental evidence to be able to serve as the host role for different human coronaviruses. Our analysis indicated that typically verified natural and laboratory human coronavirus hosts are therian mammals, esp. Eutheria mammals (or placental mammals). Second, our research identified 11 genetically engineered mouse models that were typically developed to express humanized ACE2 or DPP4 in order to become more susceptible to SARS-CoV, MERS-CoV and SARS-CoV-2 infection. From the virus side, a series of viral passages in mice were also often implemented to increase the pathogenesis/virulence of the coronaviruses to the mouse models. Third, we have modeled and represented these terms in the CIDO ontology and demonstrated its usage using SPARQL. Since currently verified human coronavirus hosts are therian mammals, it is possible that non-therian animals such as snake, turtle, chickens, and insects are not human coronavirus hosts.

To the best of our knowledge, our research provides the most comprehensive collection and classification of natural and laboratory hosts of the human coronaviruses. There have been many publications related to human coronavirus hosts, esp. COVID-19 hosts (68). However, these reports were usually focused on a specific domain of host and did not further analyze and conclude the commonness. In addition to the comprehensive collections, we focused our research on taxonomical analysis and the identification of their common taxonomical classification, resulting in the conclusion of all identified human coronavirus hosts being therian mammals. The therian mammals include placentals (Eutheria) and marsupials (Metatheria). Most of our collected hosts are placental mammals. Only one marsupial animal (i.e., *Didelphis virginiana*) was found.

Except large hairy armadillo and giant anteater, all the other natural and laboratory human coronavirus hosts under Eutheria are within Boreoeutheria, a clade of placental mammals (Figure 2 and 3). There are two broad categories under Boreoeutheria: Laurasiatheria (hedgehogs, ungulates, whales, bats, pangolins, carnivorans, and several other groups) and Euarchontoglires (rodents, lagomorphs, primates, colugos, and some other groups). In our taxonomical hierarchy, most natural animal hosts are under Laurasiatheria clade. Among them, the largest number of host animals were in the carnivora order. Unlike the natural host distribution, most laboratory animals belong to Euarchontoglires. Euarchontoglires, a superorder of placental mammals, include the following five groups: rodents, lagomorphs, treeshrews, colugos and primates. Most laboratory animals including common marmoset, African green monkey, crab eating macaque and rhesus macaque are under Simiiformes infraorder, the Primates clade.

Although all susceptible COVID-19 hosts appear to be therian mammals, not all mammals are susceptible hosts for COVID-19. The mouse model is the most frequently used laboratory animal model for infectious disease studies. While human ACE2 and murine ACE2 (mACE2) molecules are very homologous, mACE2 does not support SARS-CoV or SARS-CoV-2 binding as efficiently as hACE2 (69). To make the mouse model more useful for coronavirus study, many transgenic mouse models have been generated by expressing human ACE2 or human DPP4, both being receptors for binding to the spike protein of human coronaviruses. For example, McCray et al. reported that the transgenic expression of hACE2 in airway and other epithelia developed a rapid lethal infection after an intranasal inoculation with SARS-CoV in the transgenic mouse model (54).

Based on the results of our taxonomical classification of all the natural and laboratory animal hosts (Figure 2 and Figure 3), we hypothesize that human coronavirus hosts, esp. SARS-CoV-2 hosts are usually therian mammals. This therian host hypothesis accords with what we have known about the taxonomical classification of animal hosts of human coronaviruses. Being a scientific hypothesis, it may be proven incorrect if later contradicted observations are found. However, it is worth raising this hypothesis for now, since this hypothesis can be quite useful and it can be used to infer the non-therian mammals are not human coronavirus hosts. For example, according to the therian host hypothesis, since snake is a reptile animal under Sauropsida clade instead of a therian mammal, we then infer that snake is not a host for human coronaviruses.

Controversial data have been generated in terms of the status of the snake as a host of human coronaviruses as there has not been any detected snake that is a host of coronaviruses. A computational bioinformatics study predicted the snake to be a coronavirus potential carrier host based on its similar genetic codon usage bias with 2019-nCoV or through evolutionary analysis (e.g., SARS-CoV-2) (70, 71). However, a new study focusing on analysis of the interactions between the receptor-binding domain (RBD) of the SARS-CoV-2 S protein and the 20 key amino acid residues in the receptor ACE2 proteins from a list of mammals, birds, turtle and snake showed different results (72). Specifically, nearly half of the key residues in snake and turtles were changed. A structure simulation study showed that when a contact amino acid (AA) in hACE2 is changed to a smaller AA in snake, the contact force for the protein-protein interaction will be reduced (72). This study concluded that snake (and turtle) are not intermediate hosts for SARS-CoV-2.

To further evaluate the chance of snake being a natural host of human coronaviruses, we performed a phylogenetic tree analysis using the whole sequences of ACE2 proteins in 16 animals, including human, 13 known human coronavirus hosts, chicken (i.e., Gallus gallus), and snake (i.e., Thamnophis elegans) (Figure 5). Our results showed that the 13 ACEs from known human coronavirus hosts are phylogenetically closely related to each other, and they are distinctly far from the chicken ACE2 and even further far from the snake ACE2 (Figure 5), which confirms that snake is very unlikely an intermediate host of SARS-CoV-2.

**Figure 5.**
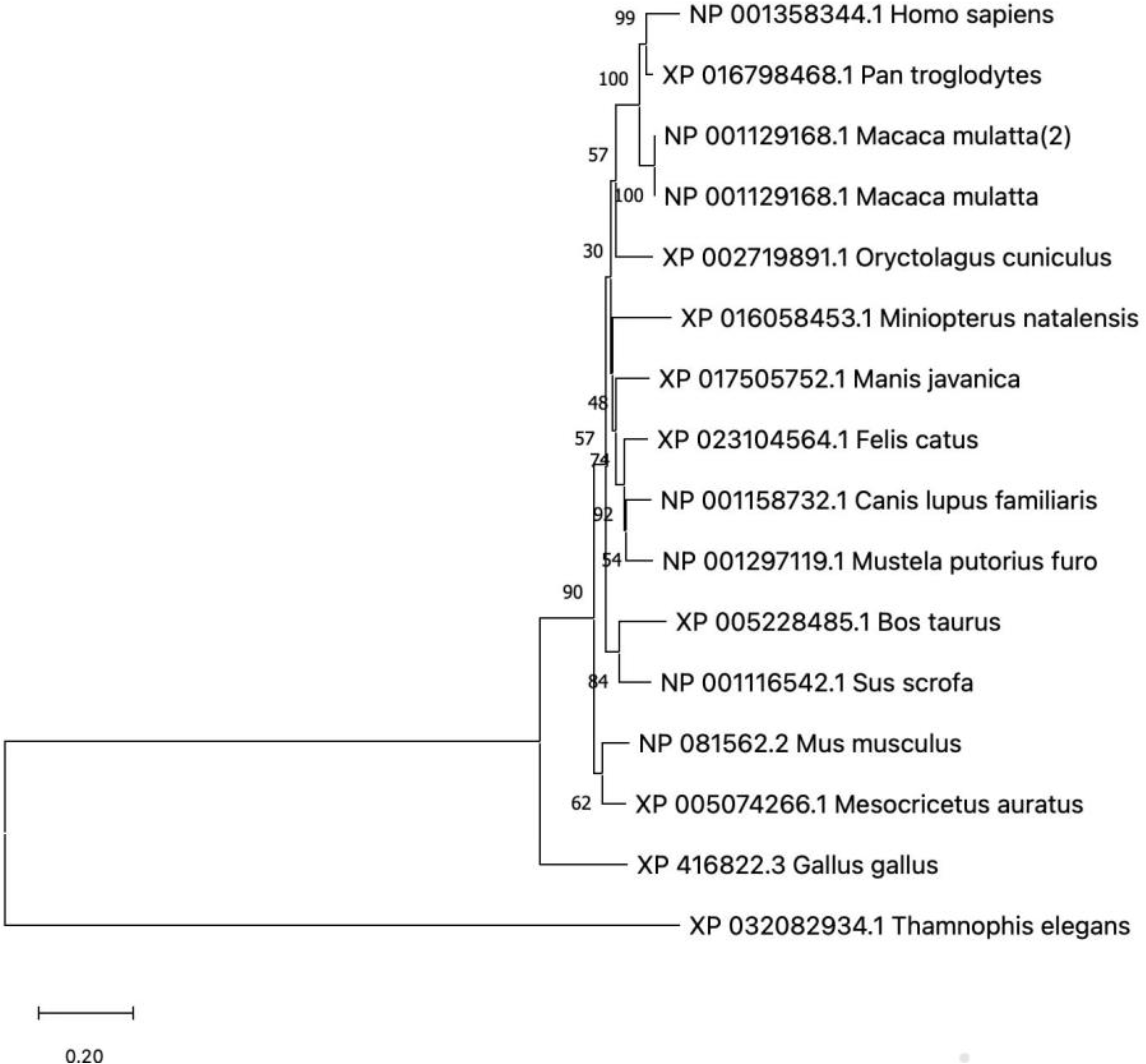
Phylogenetic analysis of ACE2 proteins from 16 animal species. The numbers represent the percentages of replicate trees in which the associated taxa were clustered together in the bootstrap test (500 replicates). The scale bar indicates nucleotide substitutions per site. The results indicate that the ACE2 proteins of *Thamnophis elegans* (i.e., Western terrestrial garter snake) and *Gallus gallus* (i.e., chicken) are phylogenetically far from those in other mammals including *Miniopterus natalensis* (i.e., Natal long-fingered bat), which aligns with our hypothesis that chicken is not susceptible to SARS-CoV-2 infection which has been previously reported (76).

Our therian animal hypothesis also excludes reptiles, birds and insects being human coronavirus hosts. Theria is a clade under the Mammalia class. Reptiles such as snakes and turtles, birds such as chickens and eagles, and insects such as housefly and mosquitoes do not belong to Mammalia or Theria. Therefore, if our hypothesis is correct, we can exclude these non-therian animals to be human coronavirus hosts. Taxonomically, there are four non-therian orders of mammals: the living Monotremata, the extinct Tricono-donta, Docodonta, and Multituberculata. Examples of the living Monotremata include Platypus and long-beaked echidnas, which all distribute in Oceania. These living Monotremata lay eggs; but like all mammals, the female montremes nurse their babies with milk. Based on our hypothesis, these living Monotremata mammals are not potential human coronavirus hosts. However, since limited data is available, more studies are required to assess our conclusions.

Based on our results and the human coronavirus therian host hypothesis proposed here, we further propose that the original host of SARS-CoV-2 would be some therian mammal. The COVID-19 pandemic was likely due to viral transmissions from nonhuman animals to human populations, leading to coronaviral evolution in nature and then infect people or through some unidentified animal host. Research evidence suggests that SARS-CoV and MERS-CoV originated in bats (73). SARS-CoV then spread from infected civets to people, while MERS-CoV spreads from infected dromedary camels to people. However, the origin of SARS-CoV-2 which caused the COVID-19 pandemic has not been identified. Our study suggests that they were likely originated from evolutions in different mammals, esp. in Boreoeutheria mammals. Furthermore, our annotation of the process of continuous coronavirus passages followed by more adaption and more virulence might imitate what happened in the gradual mutations and adaption of SARS-CoV-2 viruses in human hosts. The host-coronavirus interactions appeared to be important to the viral infection capability. Numerous studies found many genetic variations in SARS-CoV-2 and other human coronaviruses during the coronavirus evolution from their continuous interactions with the hosts (73). Such variations likely make the virus achieve “immune escape”, i.e., escaping from the host immune system and then becoming more adaptive in the host (74, 75).

Our further CIDO ontological representation makes our collection and annotations machine-interpretable in a way that computers can understand the information. Inside the CIDO ontology, such information is also seamlessly integrated with information such as protein-protein interactions and drug-target interactions. The systematical combination of all the information will allow us to better study the mechanisms of the interactions between different therian hosts and human coronaviruses. Our future work will explore how to better use the CIDO ontology for more advanced applications.

## Supporting information

Using such strategy, we have represented all the human coronaviruses and their infected hosts as identified in Tables 1-2 and Supplemental File 1.

Supplemental File 2 provides another SPARQL query that lays out the specific details of these 38 SARS-CoV-2 hosts.

## Acknowledgments

This research was supported by Youth Found of Guizhou Provincial People’s Hospital of China, GZSYQN[2019]09, the non-profit Central Research Institute Fund of Chinese Academy of Medical Sciences (2019PT320003), and a bridge fund (to YH) at the Unit for Laboratory Animal Medicine in University of Michigan Medical School.

## Author contributions

YW, XY, and YH contributed to the overall study design; YW, MY and YH provided manual verification and ontology editing; FW and HY contributed to data collection and data analysis; ZTF, HY and XY provided critical result interpretation and discussion. XY and YH secured research funds supports for YW’s research. YW and YH prepared the initial complete manuscript, and all authors contributed to the manuscript writing and reviews. All authors read and approved the final manuscript.

## Competing financial interests

The authors declare that there are no competing interests.

## Supplemental Materials

**Supplemental File 1**. Detailed lists of natural and laboratory human coronavirus hosts and transgenic mouse models. This is an Excel files with two spreadsheets, with Spreadsheet 1 providing the detailed information of natural and laboratory human coronavirus hosts, and Spreadsheet 2 providing the information of transgenic mouse models.

**Supplemental File 2**. SPARQL query of CIDO ontology for identifying specific SARS-CoV-2 hosts.

## Notes

### Competing Interest Statement

The authors have declared no competing interest.

