## Supplemental File 2 provides another SPARQL query that lays out the specific details of these 38 SARS-CoV-2 hosts. for "Taxonomical and ontological analysis of verified natural and laboratory human coronavirus hosts"

**Supplemental File 2. SPARQL query of CIDO ontology for identifying specific SARS-CoV-2 hosts**

The SPARQL script is provided below.

--

\# Goal: identify the SARS-CoV-2 hosts

PREFIX capable_of_infecting_host: <http://purl.obolibrary.org/obo/CIDO_0001195>

PREFIX SARS_CoV_2: <http://purl.obolibrary.org/obo/NCBITaxon_2697049>

SELECT distinct ?x STR(?label) as ?host_name

from <http://purl.obolibrary.org/obo/merged/CIDO>

WHERE {

?x rdf:type owl:Class .

?x rdfs:label ?label .

SARS_CoV_2: rdfs:subClassOf ?restriction .

?restriction owl:onProperty capable_of_infecting_host:; owl:someValuesFrom ?x .

}

--

**Instruction:** The following SPARQL scripts can be executed by copying the code to Ontobee SPARQL endpoint site and run: <https://ontobee.org/sparql>. See more explanation about how to run SPARQL here in our Ontobee SPARQL tutorial: <https://ontobee.org/tutorial/sparql>. More CIDO related SPARQL scripts are available here on CIDO GitHub website:

<https://github.com/CIDO-ontology/cido/blob/master/docs/sparql.txt>.

**SPARQL Output:**

The following screenshot provides the output results after executing the above SPARQL script using the Ontobee SPARQL:

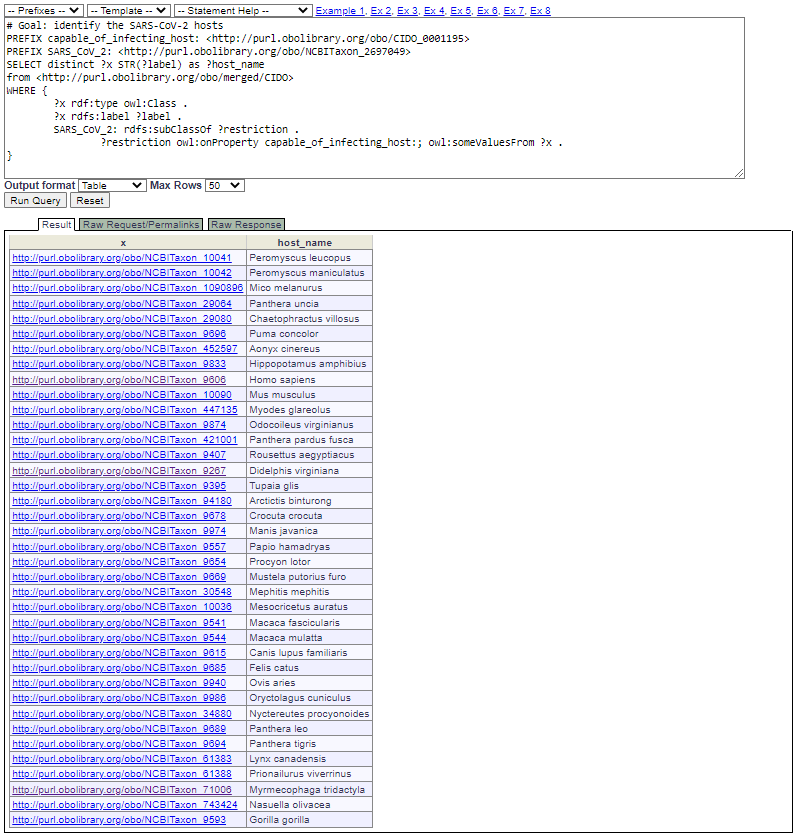

***Note:*** see more details about all human coronavirus hosts in our Supplemental File 1, which provides Excel sheets of human coronavirus hosts.
